# A Spatio-Temporal Analysis Framework for Characterizing Radiation-Induced Genomic Instability

**DOI:** 10.64898/2026.02.21.707188

**Authors:** Kriti Chopra, Clark Cucinell, Mikhail Titov, Rebecca Weinberg, Sara Forrester, Ozgur O. Kilic, Yitan Zhu, Matteo Turilli, Shantenu Jha, Daniel S. Schabacker, Thomas Brettin, Byung-Jun Yoon

## Abstract

Chronic low-dose ionizing radiation induces complex genomic instability encompassing both structural variants and point mutations, yet these alterations are typically analyzed as independent events, limiting detection of mechanistic coupling between rearrangement formation and localized mutagenesis at breakpoint junctions. This gap is particularly consequential given the widespread occupational and environmental exposure contexts such as nuclear energy, medical imaging, and environmental contamination, where coupled genomic alterations may contribute to cancer risk through mechanisms invisible to type-agnostic analyses. We developed an integrated analytical framework combining temporal pattern tracking, breakpoint-proximal mutation enrichment analysis, and systematic testing across all structural variant types to resolve these coupled dynamics across dose and time. Applying this framework to whole-genome sequencing data from primary human endothelial cells (HUVEC) exposed to chronic low-dose gamma radiation (0.20–2.62 mGy/hr) over three weeks, we identified inversion-specific mutagenic coupling; doublet base substitutions (DBS) were 7.13-fold enriched within 10bp of inversion breakpoints, a signal absent from other structural variant types, with sharp distance-dependent decay indicating localized mutagenesis at these junctions. Temporal analysis further revealed divergent fates of co-occurring alterations: inversions appeared transiently while DBS mutations showed greater persistence. These results illustrate how systematic integration of dose, time, and variant-type dimensions can uncover coupled mutagenic mechanisms that remain invisible in static or type-agnostic analyses. The framework is broadly applicable to longitudinal sequencing studies of genotoxic exposures, with applications to cancer genomics, radiation risk assessment, and mechanistic studies of DNA repair fidelity.

## 1 Introduction

Chronic low-dose ionizing radiation represents a persistent occupational and environmental hazard affecting millions of workers in nuclear energy, medical imaging, and related industries. Epidemiological studies of nuclear workers across France, the United Kingdom, and the United States have demonstrated dose-dependent increases in solid cancer mortality, with the INWORKS cohort of 309,932 workers revealing a 52% excess relative rate per Gray—estimates exceeding current radiation protection standards [Richardson et al., 2015, 2023]. Despite clear epidemiological evidence linking chronic exposure to cancer risk, the molecular mechanisms driving carcinogenesis at the genomic level remain incompletely understood, particularly under protracted low-dose exposures.

Radiation-induced genomic instability manifests as elevated rates of chromosomal aberrations, gene mutations, and cell death persisting across multiple cellular generations [Morgan, 2003a,b]. Critically, this instability occurs even at occupationally relevant exposures, with normal human fibroblasts demonstrating persistent chromosomal instability at doses as low as 100 mGy delivered at protracted dose rates [Elbakrawy et al., 2019, Khan and Wang, 2022]. Whole-genome sequencing of clonally expanded post-irradiated normal cells has further revealed that radiation induces a complex spectrum of alterations—indels, structural variants, chromothripsis, and chromoplexy—with substantial intercellular stochasticity [Youk et al., 2024]. Yet how these diverse variant classes relate mechanistically to one another remains unresolved.

The advent of whole-genome sequencing has enabled systematic cataloging of mutational signatures associated with specific mutagenic processes. The Pan-Cancer Analysis of Whole Genomes consortium identified 49 single base substitution, 11 doublet base substitution (DBS), and 17 insertion/deletion signatures across thousands of tumor genomes [Alexandrov et al., 2020], with subsequent refinement establishing rigorous methods for distinguishing true DBS signatures from coincidental adjacent substitutions [Degasperi et al., 2022]. Within this framework, radiation-specific signatures have emerged: Behjati et al. [2016] identified balanced inversions and small deletions with microhomology as hallmarks of ionizing radiation in second malignancies, while studies in mouse models demonstrated large-scale structural variants alongside oxidative stress signatures [Rose Li et al., 2020]. Localized hypermutation at rearrangement junctions is itself a recognized phenomenon: clustered substitutions co-occurring with structural variant breakpoints, termed kataegis, have been described across cancer genomes and attributed to error-prone processing of single-stranded DNA exposed at break sites [Nik-Zainal et al., 2012, Roberts et al., 2012]. Existing analytical tools, however, detect such clustering as a genome-wide, mutation-only signal, calling clustered events from inter-mutational distance within a single sample’s mutation catalog [Bergstrom et al., 2022], decoupled from the specific structural variant class and timepoint that generated it. Integrative signature models can further learn which point-mutation and structural-variant signatures co-occur across tumors [Funnell et al., 2019], but operate on aggregate persample mutation counts and likewise do not resolve coupling to individual breakpoints or timepoints. Yet whether error-prone repair couples a specific structural variant class to a specific mutation class, and how such coupling resolves over time under defined exposure, has not been systematically addressed.

The mechanistic basis for such coupling lies in error-prone double-strand break repair pathways. Cells resolve DSBs through homologous recombination, non-homologous end joining (NHEJ), microhomology mediated end joining (MMEJ), and single-strand annealing, each with distinct mutagenic potential [Ceccaldi et al., 2016]. MMEJ is inherently mutagenic, generating deletions at junctions and hypermutagenesis in flanking sequences mediated by the error-prone polymerase Pol *θ* [McVey and Lee, 2008, Seol et al., 2018, Sinha et al., 2017]. More broadly, translesion synthesis polymerases recruited to DSB junctions during NHEJ and MMEJ perform low-fidelity synthesis that can generate tandem base substitutions and complex mutations [Rodgers and McVey, 2016]. Replication-associated mechanisms such as microhomology-mediated break-induced replication add further complexity [Hastings et al., 2009]. Critically, mutational signatures are jointly shaped by initial DNA damage and subsequent repair processes, with translesion synthesis generating the majority of genotoxin-induced base substitutions [Volkova et al., 2020]. These findings suggest structural variant breakpoints—particularly those resolved by error-prone pathways—could serve as localized mutagenic hotspots, yet no systematic framework exists to test this hypothesis across variant types, doses, and time.

Existing genomic analyses of radiation exposure face a fundamental limitation: static single-timepoint sampling cannot distinguish variants arising from acute mutagenic events versus those accumulating progressively, nor can it identify which alterations persist versus those efficiently eliminated. Resolving these coupled dynamics requires integrated analysis of dose, time, and variant type. Here, we developed an integrated spatio-temporal analytical framework combining temporal pattern tracking across weekly timepoints, breakpoint-proximal mutation enrichment analysis across multiple distance windows, and systematic comparisons across all structural variant types and mutation classes. Applying this framework to whole-genome sequencing data from primary human endothelial cells exposed to chronic low-dose gamma radiation over three weeks, spanning occupationally relevant dose rates (0.20–2.62 mGy/hr) [Roach et al., 2026], we identified inversion-specific mutagenic coupling, established temporal concordance between structural variants and co-occurring point mutations, and distinguished transient versus persistent genomic damage—demonstrating the framework’s capacity to uncover coupled genomic alterations that remain invisible to type-agnostic analyses.

## 2 Results

### 2.1 Integrated framework captures structural variants and point mutations across dose and time

To investigate radiation-induced genomic alterations, we exposed primary HUVEC cells to chronic low-dose gamma radiation across five dose rates (A: 0.36, B: 0.20, C: 0.40, D: 1.47, E: 2.62 mGy/hr) [Roach et al., 2026] with matched unirradiated controls. Whole-genome sequencing was performed at weekly intervals (W1, W2, W3) following the start of exposure. We developed an integrated analytical pipeline (Figure 1) to simultaneously characterize structural variants and point mutations, with temporal pattern assignment enabling tracking of individual events across timepoints.

**Figure 1:**
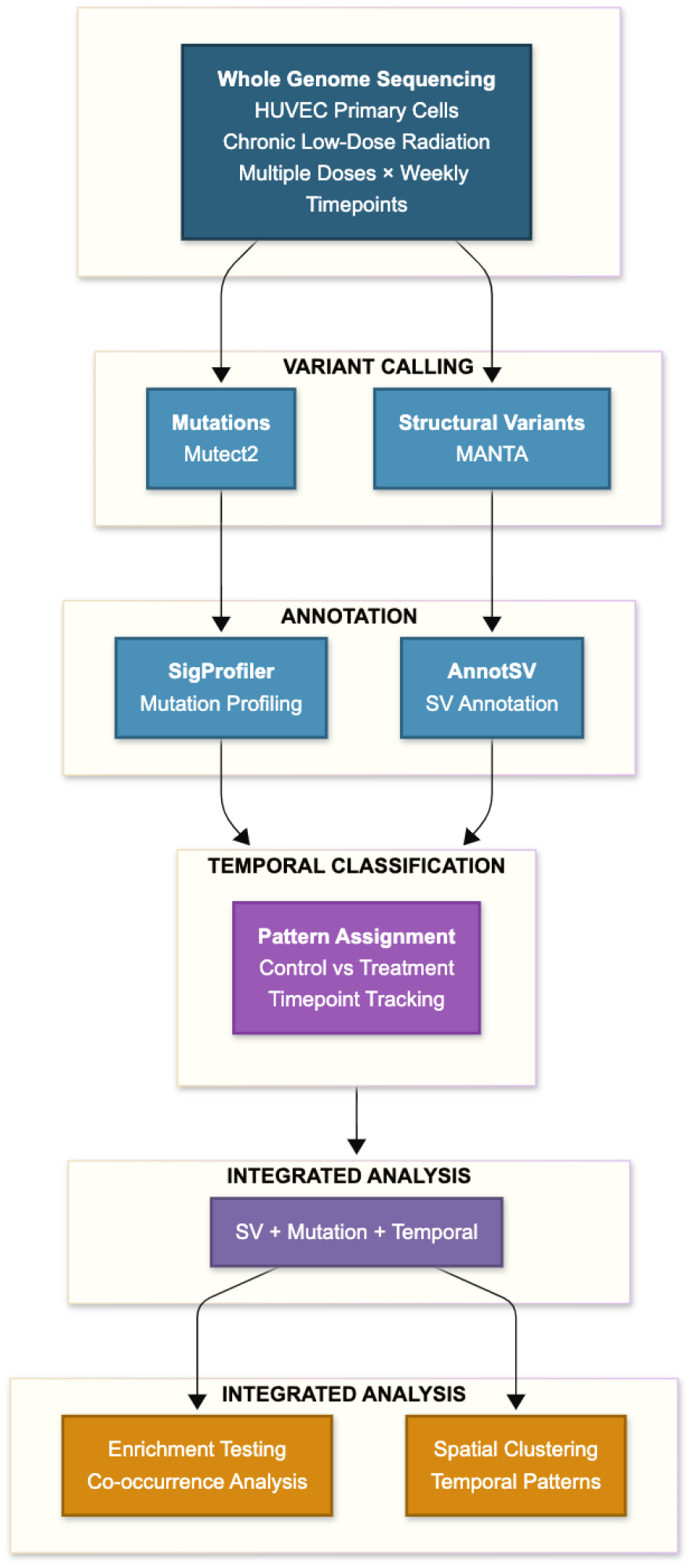
Analytical workflow for integrated structural variant and mutation analysis. Schematic overview of the computational pipeline used to analyze radiation-induced genomic alterations in HU-VEC primary cells exposed to chronic low-dose radiation across multiple dose levels(0.20–2.62 mGy/hr) and weekly timepoints (W1–W3). Whole genome sequencing data was processed through parallel variant calling pipelines: Mutect2 for point mutations and MANTA for structural variants (SVs). Mutations were profiled using SigProfiler to extract mutational signatures, while SVs were annotated using AnnotSV for functional characterization. Temporal classification assigned pattern codes to each variant based on presence/absence across timepoints in treatment versus control samples. The integrated analysis combined SV coordinates, mutation positions, and temporal patterns to perform enrichment testing, co-occurrence analysis, spatial clustering, and temporal pattern characterization.

Structural variants were called using MANTA and annotated with AnnotSV, while point mutations were identified using Mutect2 and profiled with SigProfiler. Each variant was assigned a temporal pattern code based on presence/absence across timepoints in treatment versus control samples (e.g., “0T00” for variants appearing only at week 1 in treated samples, “0TTT” for persistent variants present at all three timepoints). This integrated framework characterized all four mutation classes (single nucleotide variants [SNV], doublet base substitutions [DBS], multi-nucleotide substitutions [MNS], and insertions/deletions [InDel]) alongside five structural variant types (translocations [TRA], inversions [INV], duplications [DUP], deletions [DEL], and insertions [INS]) across 19 samples, enabling dose- and time-resolved analysis of radiation-induced genomic alterations.

### 2.2 Temporal pattern encoding resolves progressive mutation accumulation across the exposure period

We examined the temporal dynamics of mutation accumulation across the radiation exposure period. Sankey analysis was performed for all mutation types; results for DBS are shown as the primary mutation class of interest (Figure 2). The dominant flow at each timepoint was the Both category, i.e. mutations detected in both irradiated and control samples, reflecting a shared mutational background between treated and untreated cells. Radiation exposure additionally generated a continuous stream of new Exposed mutations (detected only in irradiated samples) at each timepoint, while the control arm showed consistently smaller and stable flows, indicating that radiation substantially amplifies mutation induction above the background level of spontaneous mutagenesis. This parallel tracking of control-arm mutations provides a direct baseline for distinguishing radiation-induced from replication-associated mutagenesis inherent to normal cell proliferation.

**Figure 2:**
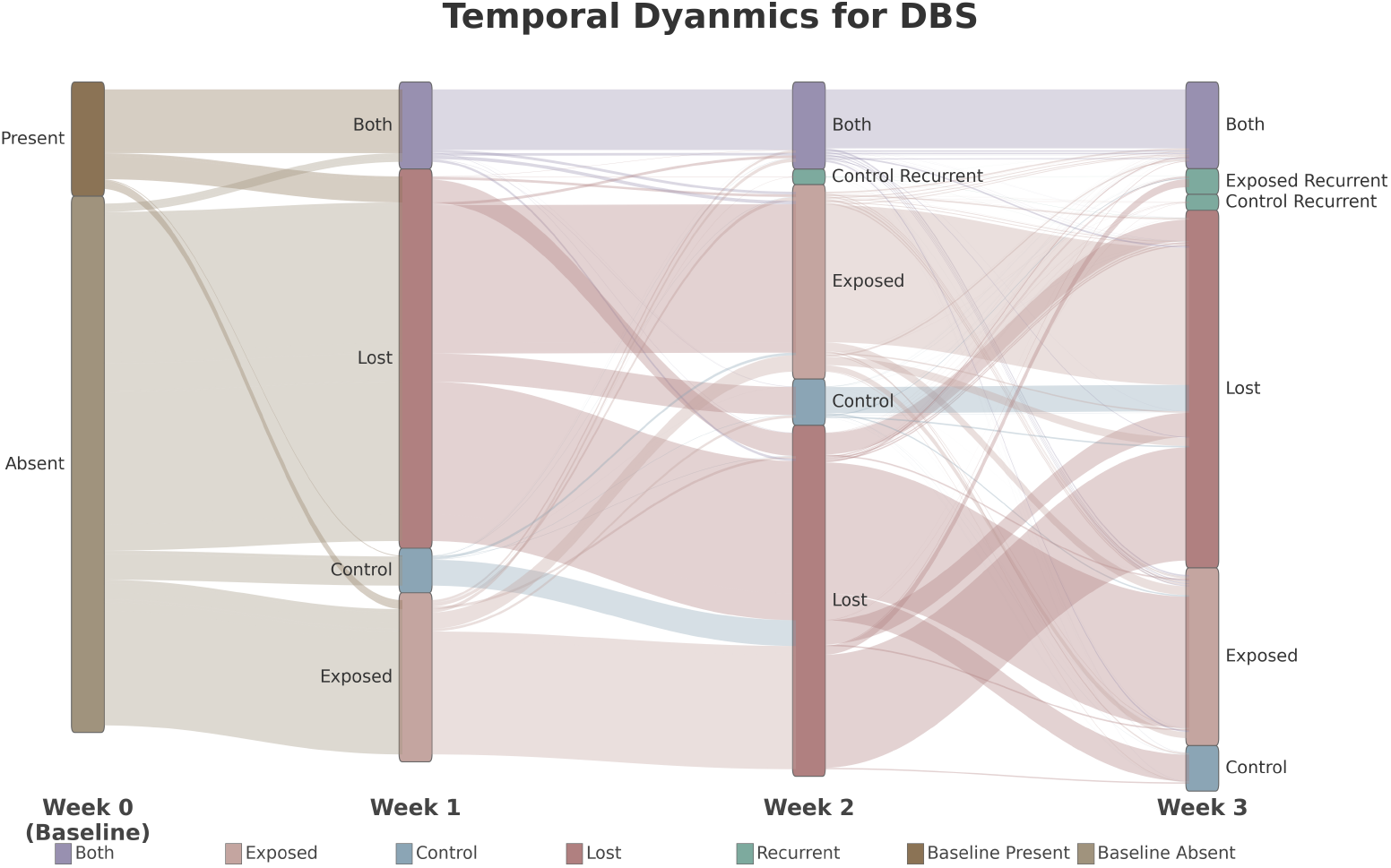
Temporal dynamics of doublet base substitution accumulation. Sankey diagram illustrating DBS mutation flow across weekly timepoints (W0–W3), aggregated across all radiation dose levels and chromosomes. Flow width is proportional to mutation count. At baseline (Week 0), mutations are classified as present (brown) or absent (gray). Present mutations either persist through subsequent timepoints (purple flows to “Both”) or are cleared from the population (dusty rose flows to “Lost”). New mutations emerge from the baseline-absent pool and are classified as radiation-induced (“Exposed”, pink) if detected only in irradiated samples, or spontaneous (“Control”, blue) if detected in control samples. A subset of mutations classified as “Recurrent” (teal) reappear at Week 3 after being absent at intermediate timepoints. The progressive accumulation of exposed mutations from W1 to W3 demonstrates radiation-dependent mutation induction over the chronic exposure period.

Mutations present at baseline either persisted through subsequent timepoints or were cleared from the population, while new mutations emerged predominantly in irradiated samples. A subset of mutations showed multi-timepoint persistence (“0TTT” pattern), suggesting clonal expansion of mutation-carrying cells. Small but visible Exposed Recurrent and Control Recurrent flows at weeks 2 and 3 indicate that a fraction of mutations reappeared after prior absence—for example, mutations with pattern “0T0T” reappearing at W3 after absence at W2, or “00TT” appearing first at W2 and persisting to W3—reflecting complex clonal dynamics or detection threshold effects at the sequencing depth used. Qualitatively similar temporal dynamics were observed across SNV, MNS, and InDel mutation classes, with SNV and InDel showing larger baseline-present contributions relative to MNS (Figure S1). Quantitative counts of all seven radiation-specific temporal patterns across mutation types are provided in Table S1.

### 2.3 Framework characterizes the structural variant landscape across dose and time

Characterization of the SV landscape (Figure 3) revealed translocations (TRA) as the predominant SV type (78,428 unique events), followed by inversions (INV; 8,716), duplications (DUP; 1,367), deletions (DEL; 801), and insertions (INS; 3). TRA events dominated across all dose levels, consistent with radiation-induced inter-chromosomal rearrangements [Youk et al., 2024].

**Figure 3:**
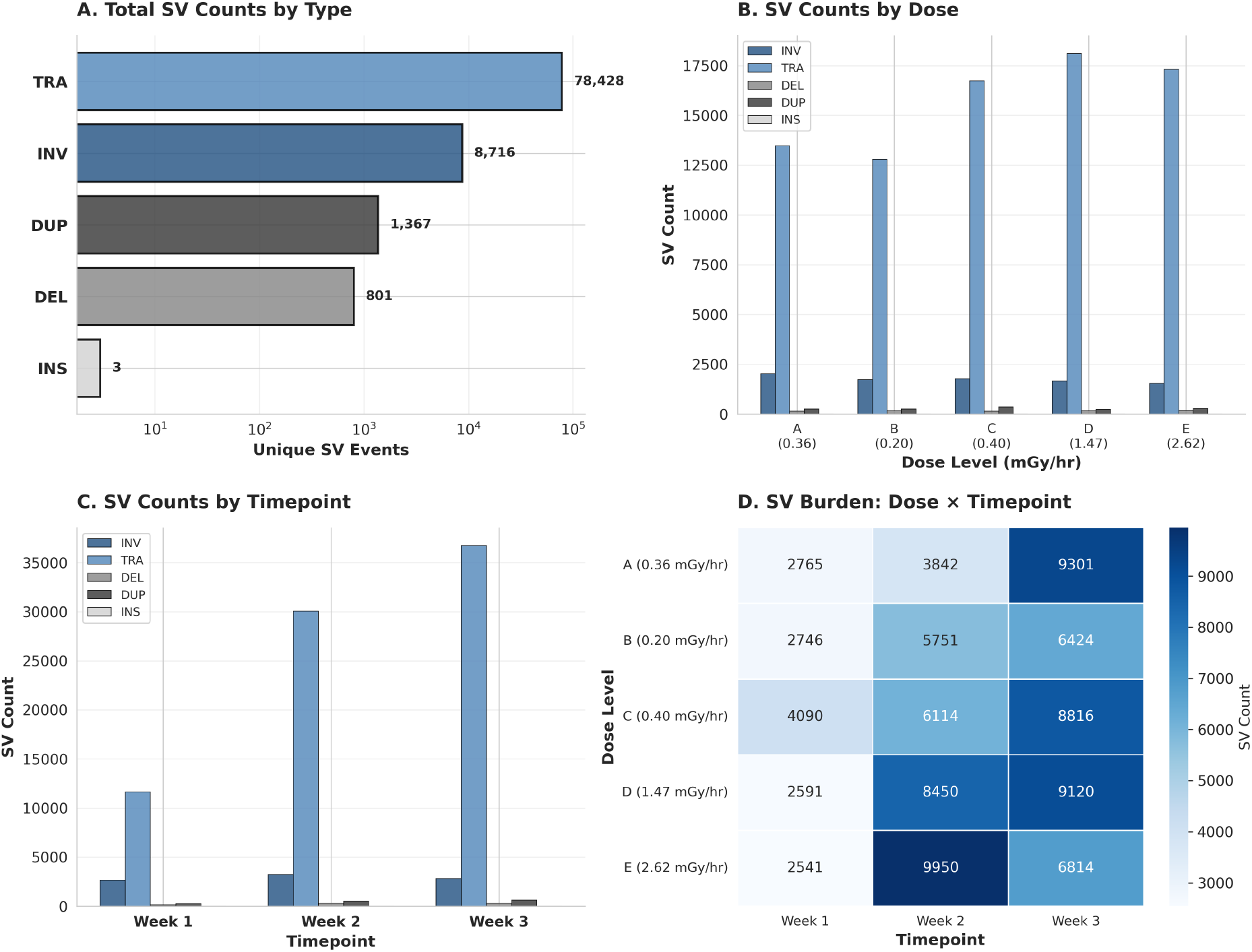
Structural variant landscape across dose and time. **(A)** Total unique SV events by type (log scale). Translocations (TRA) predominate (78,428), followed by inversions (INV, 8,716), duplications (DUP, 1,367), deletions (DEL, 801), and insertions (INS, 3). **(B)** SV counts by dose level (A: 0.36, B: 0.20, C: 0.40, D: 1.47, E: 2.62 mGy/hr), grouped by SV type. TRA events dominate across all doses. **(C)** SV counts across timepoints. Total SV burden increases from Week 1 to Week 3, driven primarily by TRA. **(D)** Heatmap of total SV burden by dose and timepoint. All doses show temporal accumulation, with highest counts at Week 3 except that dose E (2.62 mGy/hr) shows peak burden at Week 2.

Temporal analysis demonstrated progressive SV accumulation, with total burden increasing from Week 1 to Week 3 across all dose levels. The dose-timepoint heatmap (Figure 3D) revealed that while all doses showed temporal accumulation, the highest dose (E: 2.62 mGy/hr) exhibited a distinctive pattern with peak burden at Week 2 followed by slight reduction at Week 3, potentially reflecting dose-dependent cellular elimination mechanisms. Temporal pattern analysis revealed that SVs across all types were predominantly single-timepoint events (0T00, 00T0, 000T), with W3-dominant patterns (000T) comprising the largest fraction (46.8% across all doses). Notably, no SV of any type showed persistence across all three timepoints (0TTT pattern), in striking contrast to point mutations which accumulate progressively across the exposure period—a divergence consistent with preferential cellular elimination of structural aberrations relative to point mutations.

Analysis of repetitive element involvement (Figure S2) showed that 61.7% of SVs had both breakpoints within interspersed repeat elements, with LINE and Alu elements predominating. Radiation-induced SVs showed slightly lower repeat involvement (66.2%) compared to controls (69.8%, Δ = −3.6%), suggesting that radiation may induce breakpoints in non-repetitive regions at modestly higher rates. Family-level distributions were largely similar between conditions (Table S2), with the notable exception of Alu elements, which were proportionally underrepresented at radiation-induced breakpoints (20.3% vs 25.7% in controls, Δ = −5.4 pp). This Alu depletion accounts for the majority of the overall reduction in repeat involvement under radiation exposure.

### 2.4 Type-stratified enrichment analysis identifies inversion-specific DBS coupling

Building on the dose- and time-resolved catalog of structural variants and point mutations established by our integrated framework, we next investigated whether SV breakpoints exhibit localized enrichment of point mutations, a pattern that would implicate error-prone repair as a secondary mutagenic mechanism operating at sites of structural rearrangement. We counted mutations within defined distance windows of breakpoints across all SV events, comparing radiation-induced versus control mutation frequencies (Table S3). At the smallest window (10bp), radiation-induced DBS showed significant enrichment at SV breakpoints (1.55×, Poisson test, *P* = 1.65 × 10^*−*12^), while control DBS showed no enrichment (0.96 ×, *P* = 0.74). This radiation-specific signal decayed with distance, losing statistical significance beyond 25bp (*P* = 4.59 × 10^*−*4^ at 25bp; *P* = 0.89 at 50bp). Among other mutation types, SNV and MNS showed modest radiation-specific enrichment at 10bp (1.24× and 1.30×, respectively), while InDel mutations showed comparable enrichment in both radiation and control samples across all window sizes (Table S3), consistent with indel generation as a general feature of double-strand break repair rather than a radiation-specific mutagenic process [Ceccaldi et al., 2016].

To identify which SV types drive this signal, we stratified enrichment analysis by structural variant type (Figure 4). Strikingly, inversion breakpoints showed 7.13-fold enrichment of DBS within 10bp windows compared to control mutations, a signal absent from other SV types. Translocations showed only 1.55× enrichment, duplications 1.22×, and deletions 0.29× (below baseline). The specificity of this signal demonstrates that the mutagenic coupling is intrinsic to the inversion formation mechanism rather than a general property of structural rearrangements. Analysis across all mutation types at inversion breakpoints revealed that DBS showed the strongest coupling (7.13× ), with moderate enrichment for multi-nucleotide substitutions (MNS, 1.64× ), while single nucleotide variants (SNV, 1.41×) and insertions/deletions (InDel, 0.85× ) showed minimal or no enrichment. The predominance of DBS suggests error-prone translesion synthesis operating specifically at inversion junctions, where tandem base misincorporations occur during repair of complex DNA structures [Rodgers and McVey, 2016, Seol et al., 2018].

**Figure 4:**
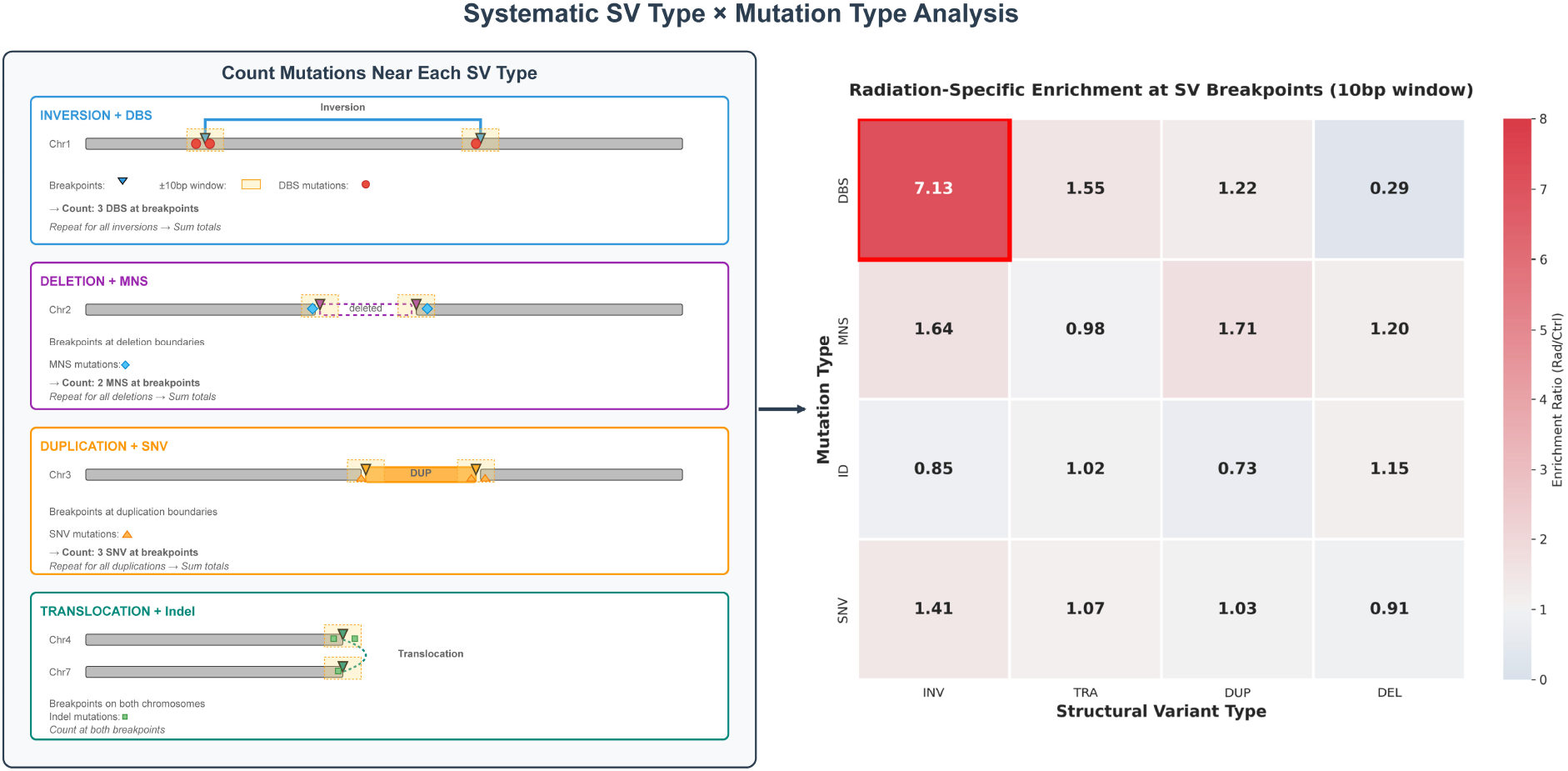
Mutation type and structural variant coupling specificity. **(Left)** Schematic of the analytical approach: for each SV type, mutations within ±10bp of breakpoints are counted across all events, comparing radiation-induced and control mutation frequencies. **(Right)** Heatmap showing radiation-specific enrichment (radiation/control ratio) for each mutation type (rows) at breakpoints of each SV type (columns) within 10bp windows. Doublet base substitutions (DBS) at inversion (INV) breakpoints show striking enrichment (7.13 ×, red box), substantially exceeding all other mutation-SV combinations. Moderate enrichment is observed for multi-nucleotide substitutions (MNS) at INV (1.64× ) and DUP (1.71× ) breakpoints. Single nucleotide variants (SNV) and insertions/deletions (InDel) show minimal enrichment across all SV types (0.73–1.41 ×). Values represent the ratio of observed/expected enrichment in radiation samples divided by the same ratio in control samples.

The distance-dependent decay analysis (Figure S3 A) revealed that INV-DBS coupling shows steep decay from 7.13× at 10bp to approximately 1.9× at 100bp, while other SV types (TRA, DUP, DEL) showed minimal or no enrichment across all distances. When examining different mutation types at inversion breakpoints (Figure S3 B), DBS exhibited strong distance-dependent decay (7.13× to ∼1.8 ×), while MNS, SNV, and InDel remained near baseline (0.8–1.6 ×) regardless of distance. This sharp spatial decay demonstrates that the mutagenic effect is tightly localized to the immediate breakpoint junction, consistent with error-prone repair during inversion formation rather than regional chromatin effects or replication stress extending across broader genomic domains.

To test whether spatially proximal INV-DBS pairs arise from shared mutagenic processes rather than independent events at the same locus, we performed temporal concordance analysis on 48 co-occurring pairs within 10bp (Figure S4),of which 41 (85.4%) were temporally concordant. Inversions and DBS mutations showed similar temporal accumulation patterns across timepoints (W1: 30.6% vs 27.6%, W2: 37.1% vs 32.2%, W3: 32.3% vs 31.2%), but differed dramatically in persistence: inversions were exclusively transient (100% single-timepoint), while DBS showed modest multi-timepoint persistence (9.0%). Critically, temporal concordance defined as INV-DBS pairs sharing at least one common timepoint substantially exceeded random expectation (33%) at all timepoints: W1 (66.7%, *χ*^2^ = 6.00, *P* = 1.43 ×10^*−*2^), W2 (85.0%, *χ*^2^ = 24.02, *P* = 9.51 ×10^*−*7^), and W3 (100%, *χ*^2^ = 32.00, *P* = 1.54 ×10^*−*8^), with overall concordance of 85.4% (*χ*^2^ = 58.59, *P* = 1.94 ×10^*−*14^). The temporal locking was tight: all concordant DBS mutations were single-timepoint and matched the inversion time-point exactly, with no concordant pairs arising from persistent DBS patterns that incidentally included the inversion’s timepoint. This high concordance suggests that co-localized inversions and DBS mutations likely arise from shared mutagenic processes rather than independent events at the same locus. Concordance analysis by inversion size revealed that small (*<*10kb, n=42), medium (10kb–1Mb, n=2), and large (1–50Mb, n=2) inversions all showed high concordance (88–100%), while mega-inversions ( ≥50Mb, n=2) showed none; the coupling between inversion formation and DBS mutagenesis thus operates across inversion sizes up to the megabase scale.

### 2.5 Dose-stratified analysis reveals scaling of functional impact through inversion size

Analysis of INV-DBS events by radiation dose (Figure 5) revealed striking differences in genomic distribution and functional impact. Of the 41 temporally concordant INV-DBS pairs, low-dose exposure (0.20–0.40 mGy/hr; groups A,B,C) and high-dose exposure (1.47–2.62 mGy/hr; groups D,E) generated similar numbers (20 vs 21), but substantially different numbers of affected genes (5 vs 52). Note that the 20 low-dose INV-DBS pairs shown in Figure 5B represent individual pair counts, whereas Figure 5A displays the genomic loci of the inversions involved: multiple pairs mapping to the same or overlapping genomic positions are represented as a single wider bar, resulting in fewer visible loci (11 bars) than total pairs. Furthermore, the 11 loci span 9 chromosomes yet affect only 5 protein-coding genes because the majority of inversion events fall in non-coding or intergenic regions; only inversions whose annotated gene overlap (from AnnotSV) includes a protein-coding entry contribute to the gene count. The genome-wide distribution (Figure 5A) showed that both dose groups produced scattered INV-DBS events across multiple chromosomes. Low-dose events consisted primarily of small inversions (*<*1Mb, thin lines), while high-dose events included both small and large inversions (≥ 1Mb, thicker lines). The presence of large inversions at high doses contributed to the greater gene impact observed in the high-dose group.

**Figure 5:**
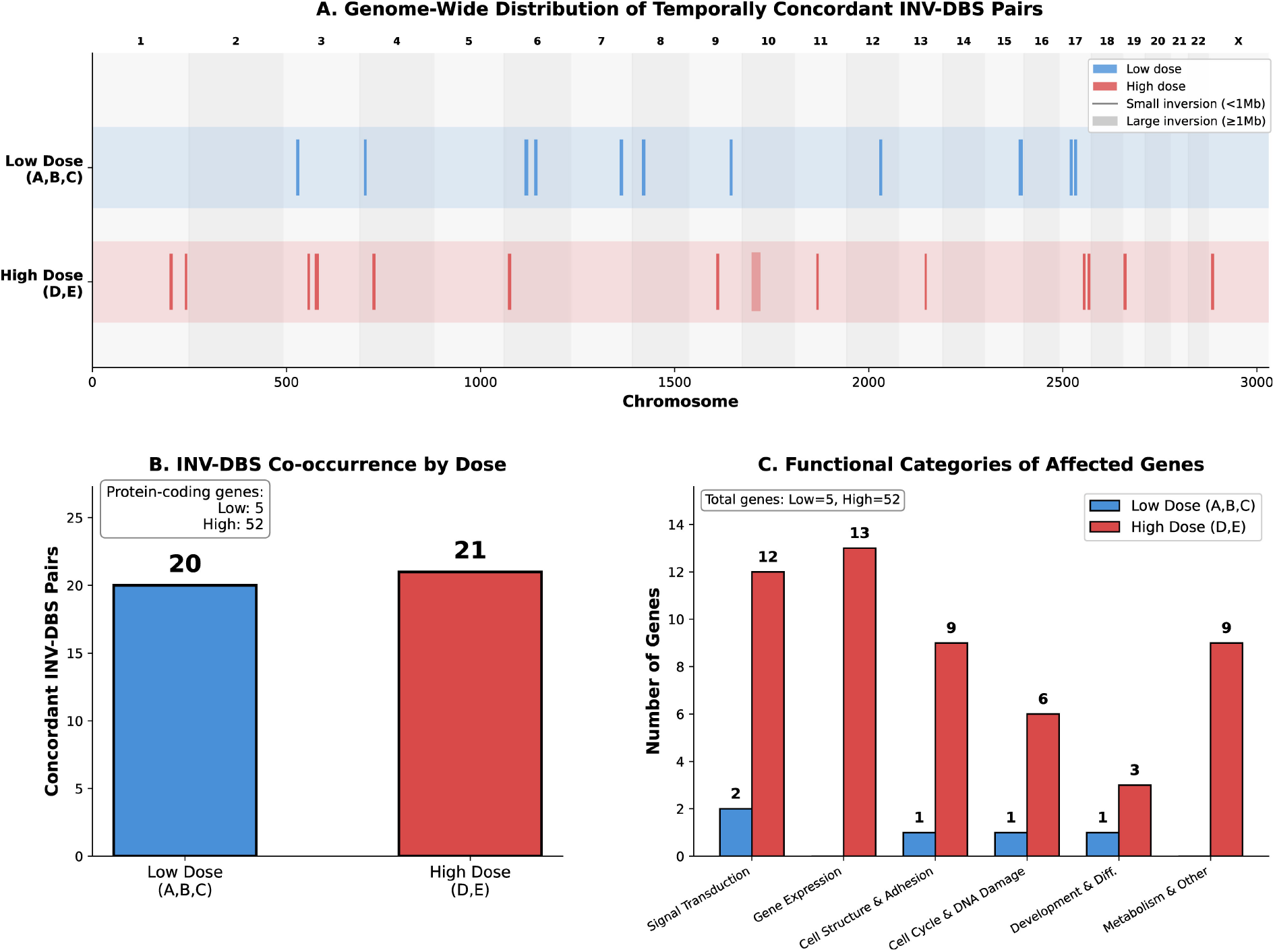
Dose-dependent genomic distribution and functional impact of INV-DBS events. **(A)** Genome-wide distribution of INV-DBS co-occurrence events by dose group. Low dose (A,B,C: 0.20–0.40 mGy/hr, blue) and high dose (D,E: 1.47–2.62 mGy/hr, red) groups show distinct patterns. Vertical lines indicate affected loci; line thickness represents inversion size (thin: *<*1Mb small inversions, thick: ≥1Mb large inversions). High-dose events include more large inversions distributed across multiple chromosomes. **(B)** INV-DBS pair counts by dose group. Similar event counts (Low: 20, High: 21) mask substantial differences in gene impact (Low: 5 genes, High: 52 genes), reflecting large-inversion involvement at high doses. **(C)** Functional categories of INV-DBS affected genes. High-dose events (red) predominantly affect gene expression (13), signal transduction (12), cell structure and adhesion (9), cell cycle and DNA damage (6), development and differentiation (3), and metabolism and other functions (9). Low-dose events (blue) show minimal gene involvement across categories, with signal transduction (2), cell structure and adhesion (1), cell cycle and DNA damage (1), and development and differentiation (1) affected.

Functional categorization (Figure 5C) revealed that high-dose INV-DBS events affected genes involved in gene expression (13 genes), signal transduction (12), cell structure and adhesion (9), cell cycle and DNA damage (6), metabolism and other functions (9), and development and differentiation (3). Low-dose events showed minimal gene involvement, with signal transduction (2), cell structure and adhesion (1), cell cycle and DNA damage (1), and development and differentiation (1) being the only affected categories. Given the stochastic nature of radiation-induced structural variants, the greater functional impact at higher doses reflects the increased probability of large inversions spanning gene-dense regions rather than systematic dose-dependent pathway targeting. Nonetheless, the involvement of 16 high-constraint loci (pLI ≥0.9) (Table S4), among concordant INV-DBS events, including genes functioning in DNA damage response and signal transduction, highlights the potential functional consequences when such coupled lesions do occur.

## 3 Discussion

The central contribution of this study is an integrated analytical framework that combines temporal pattern tracking, breakpoint-proximal enrichment analysis, and structural variant type-specific comparisons to resolve when, where, and in what form coupled genomic alterations arise following radiation exposure. This addresses a fundamental gap in radiation genomics: while individual variant classes have been extensively cataloged, their mechanistic coupling at breakpoint junctions has remained inaccessible to type-agnostic analyses. Applying the framework to chronic low-dose gamma radiation in primary human endothelial cells, we discovered that inversion breakpoints exhibit 7.13-fold enrichment of doublet base substitutions within 10bp of junctions, a signal absent from other structural variant types. This finding demonstrates how systematic integration of dose, time, and variant-type dimensions can uncover coupled mutagenic patterns that remain invisible in static or single-class analyses.

The specificity of INV-DBS coupling restricted to inversions among all SV types and to DBS among all mutation types points to error-prone DNA repair as a candidate mechanism. The sharp distance-dependent decay (7.13× at 10bp to ∼1.9× at 100bp) is consistent with localized action of translesion polymerases at breakpoint junctions, likely operating through microhomology-mediated end joining (MMEJ) or Pol *θ*-mediated pathways [Rodgers and McVey, 2016, Seol et al., 2018]. Unlike deletions or duplications, inversions require re-ligation of DNA ends in inverted orientation, a topology that may preferentially engage error-prone repair mechanisms when canonical NHEJ cannot efficiently resolve strand polarity mismatches [Porubsky et al., 2022]. This suggests inversions function not as isolated structural events but as compound lesions that couple rearrangement formation with clustered point mutation, plausibly through a shared error-prone repair process. Notably, the INV-DBS events identified here were uniformly isolated single doublet substitutions at inversion breakpoints, with one DBS per junction across all concordant pairs, rather than the multi-substitution tracts that define kataegis [Nik-Zainal et al., 2012, Roberts et al., 2012]; this single-event multiplicity, together with the doublet rather than single-base character of the substitutions, points to a discrete error-prone repair event at the junction rather than processive deamination of single-stranded DNA.

The temporal dimension of our framework proved essential for evaluating whether co-localized lesions share a mechanistic origin. Temporal concordance between spatially co-occurring INV-DBS pairs (85% overall) substantially exceeded the 33% random expectation and rose across the exposure period, suggesting that these lesions arise from shared mutagenic processes rather than coincidental spatial overlap. The contrasting fates of the two variant classes, transient inversions versus more persistent DBS, are consistent with differential cellular quality control that preferentially eliminates structural aberrations while retaining point mutations. This interpretation aligns with recent evidence that DNA repair leaves lasting genomic marks beyond sequence restoration: even successfully repaired loci exhibit heritable chromatin alterations and transcriptional impairment [Bantele et al., 2025], suggesting that cells cleared of structural variants may retain both sequence-level mutations and epigenomic dysfunction at former breakpoint sites. This divergence holds across the broader SV landscape and likely reflects the greater fitness cost of chromosomal rearrangements relative to point mutations.

The dose-response analysis revealed a decoupling between event frequency and functional impact: similar numbers of INV-DBS events at low and high doses produced a roughly 10-fold difference in affected genes (5 vs 52). Because radiation-induced structural variants arise stochastically, this difference reflects the greater genomic reach of the larger inversions occurring at higher dose rates rather than systematic pathway targeting. Nonetheless, among the 16 high-constraint genes (pLI ≥0.9) affected by temporally concordant events, several function in DNA damage response (WAC, CUL2, EPC1), highlighting the potential functional consequences when such coupled lesions overlap constrained loci.

Beyond the specific findings reported here, the spatio-temporal framework applies to other longitudinal sequencing contexts in which coupled genomic alterations may operate. The temporal pattern encoding system, which tracks individual variants across timepoints through a four-position presence/absence classification in treatment versus control samples, generalizes to any longitudinal cohort with matched controls, with potential applications to chemotherapy-induced mutagenesis studies, aging-associated genomic instability cohorts, and cancer evolution datasets with serial sampling. The breakpoint-proximal enrichment analysis, stratified by SV type, can be applied to any genotoxic exposure to test whether specific rearrangement classes generate coupled point mutations. Together, temporal tracking, type-stratified enrichment, and concordance testing provide a unified analytical approach for investigating relationships between variant classes that have traditionally been analyzed independently.

Several methodological considerations warrant discussion. First, our temporal resolution (weekly timepoints) cannot distinguish variants arising within the same cell cycle versus across successive divisions, though the temporal concordance analysis mitigates this by requiring dose- and timepoint-matched comparisons. Second, while the enrichment pattern is consistent with MMEJ/Pol *θ* pathways, direct mechanistic validation would require genetic perturbation using polymerase-deficient cell lines or specific inhibitors. Third, the predominance of large inversions at higher dose rates means that stochastic large-scale events disproportionately influence gene-level associations, reflecting the genomic reach of individual structural variants rather than targeted mutagenic hotspots. Fourth, like all short-read SV analyses, our findings depend on the specific calling algorithm used (MANTA with AnnotSV-based reclassification of paired breakend records into inversions). Different SV callers vary in their inversion call sets due to algorithmic differences in handling paired breakends and complex repair junctions; orthogonal validation with long-read sequencing or breakpoint-spanning PCR would be required to further characterize individual inversion events. Finally, bulk whole-genome sequencing cannot resolve subclonal structure, meaning that temporally concordant INV-DBS pairs detected at the same dose and timepoint are likely but not guaranteed to arise within the same cell or subclone. Our focus on spatially proximal, temporally matched pairs enriches for events sharing a common mutagenic origin, though definitive single-cell co-occurrence would require read-level overlap of both variants or single-cell sequencing approaches. Despite these constraints, the compound nature of inversion lesions indicates that analyses treating structural variants and point mutations independently may underestimate the mutagenic impact of radiation. Consistent with evidence that DSB repair itself generates localized mutations [Bader and Bushell, 2023], this coupling may represent a specific instance of a general phenomenon in which repair fidelity costs extend mutagenic consequences beyond the primary lesion.

In conclusion, our integrated spatio-temporal framework reveals radiation-induced mutagenesis as a multi-dimensional process in which structural variant type, genomic location, dose level, and temporal dynamics jointly determine functional outcomes. By combining temporal pattern tracking, breakpoint-proximal enrichment analysis, and SV type-specific comparisons, all unified through a four-position encoding that tracks individual variants across dose and time, the framework provides a generalizable approach for distinguishing transient from persistent alterations and investigating coupled genomic damage in any longitudinal sequencing study, with applications spanning radiation risk assessment, cancer genomics, and mechanistic studies of DNA repair fidelity. As a demonstration of its utility, the framework identified inversion breakpoints as sites of localized secondary mutagenesis under chronic low-dose gamma radiation, with affected high-constraint genes functioning in DNA damage response and signal transduction pathways critical for maintaining genomic stability.

## 4 Materials and Methods

### 4.1 Cell Culture and Radiation Exposure

Cell culture and radiation exposure methods are described in detail in [Weinberg et al., 2026] and [Roach et al., 2026]. Briefly, primary human umbilical vein endothelial cells (HUVEC; PromoCell, Cat. No. C-12200) were cultured in Human Large Vessel Endothelial Cell Basal Medium (Gibco, Cat. No. M-200-500) supplemented with 1X Large Vessel Endothelial Supplement (Gibco, Cat. No. A1460801). Cells were maintained in a humidified incubator at 37°C with 5% CO_2_ and passaged at 70–90% confluency.

For radiation exposure, cells were cultured in custom 96-well source plate configurations containing ^137^Cs sources, as detailed in [Roach et al., 2026]. Cells were exposed to five dose rates: experimentally determined as 0.36 mGy/hr (Group A), 0.20 mGy/hr (Group B), 0.40 mGy/hr (Group C), 1.47 mGy/hr (Group D), and 2.62 mGy/hr (Group E) [Roach et al., 2026], corresponding to cumulative doses of approximately 181, 101, 202, 741, and 1,321 mGy over three weeks, respectively. Matched unirradiated controls were maintained in a separate incubator under identical culture conditions. Both irradiated and matched unirradiated control cells were harvested at weekly intervals (W0 baseline, W1, W2, W3) for nucleic acid extraction, yielding a total of 19 DNA sequencing samples (5 dose levels ×3 timepoints for irradiated cells, plus 4 timepoints for unirradiated controls).

### 4.2 Whole Genome Sequencing

DNA extraction, library preparation, and sequencing methods are described in detail in [Weinberg et al., 2026]. Briefly, shotgun genomic DNA libraries were constructed from 50 ng of DNA using the Illumina DNA Prep kit with unique dual-indexed adaptors to prevent index switching. Library construction and sequencing were performed at the Roy J. Carver Biotechnology Center, University of Illinois at Urbana-Champaign. Individually barcoded libraries were quantified using a Qubit fluorometer (Thermo Fisher) and evaluated on a Fragment Analyzer (Agilent) to verify the absence of primer dimers and confirm expected fragment sizes. Libraries were pooled at equimolar concentration and the pool was further quantified by qPCR on a BioRad CFX Connect Real-Time System (BioRad Laboratories). Pooled libraries were loaded across two 25B lanes of a NovaSeq X Plus (Illumina) and sequenced with paired-end reads of 150 bp from each end. Raw sequencing reads were demultiplexed using bcl2fastq v2.20 and quality-assessed with FastQC software.

### 4.3 Point Mutation Calling and Classification

Bioinformatics workflows for variant calling are described in detail in [Weinberg et al., 2026]. Briefly, raw FASTQ files were assessed for quality using FastQC and trimmed using TrimGalore (v0.6.10) to remove adapters and low-quality bases. High-quality reads were aligned to the GRCh38/hg38 human reference genome using minimap2 (v2.28-r1209) in short-read mode (`-ax sr`). Following the GATK Somatic Short Variant Discovery best practices workflow, somatic variants were called using Mutect2 (GATK v4.5.0.0) [Benjamin et al., 2019] with matched untreated controls across 50 scattered genomic intervals in parallel, and resulting interval VCFs were merged using GATK GatherVcfs. Cross-sample contamination was estimated using GATK GetPileupSummaries and CalculateContamination prior to filtering. Variants were filtered using FilterMutectCalls with default parameters. The resulting VCF files were annotated using GATK Funcotator (v1.8) in MAF format against the hg38 reference.

Mutation classification into single nucleotide variants (SNV), doublet base substitutions (DBS), multi-nucleotide substitutions (MNS), and insertions/deletions (InDel) was performed using SigProfiler-MatrixGenerator [Bergstrom et al., 2019] and SigProfilerExtractor [Islam et al., 2022] with the GRCh38 reference genome. SigProfiler output files were parsed to extract mutation coordinates, alleles, and sample information. Each mutation was assigned a unique identifier based on genomic position and sample to enable tracking across timepoints. Mutations were annotated with overlapping genomic features using UCSC refGene annotations (GRCh38).

### 4.4 Structural Variant Calling

Structural variants (SVs) were called using MANTA v1.6.0 [Chen et al., 2016] with default parameters for somatic variant calling. Tumor-normal pairs consisted of irradiated samples (tumor) versus matched baseline (W0) samples (normal). SVs passing MANTA’s internal quality filters (FILTER = PASS) were retained for downstream analysis.

SV annotation was performed using AnnotSV v3.3 [Geoffroy et al., 2018]. SVs were classified by type (inversions, translocations, deletions, duplications, insertions) based on MANTA’s SVTYPE field. AnnotSV reclassified a subset of breakend (BND) calls as inversions based on paired breakpoint orientation and MATEID annotations, consolidating fragmented inversion calls into single events. Annotations included gene overlap, repeat element involvement, and functional impact predictions.

### 4.5 Temporal Pattern Assignment

Each variant was assigned a temporal pattern code based on presence/absence across the four timepoints (W0, W1, W2, W3) and treatment condition (radiation vs control). Pattern codes use four positions where “0” indicates absence and “T” (irradiated) or “C” (control) indicates presence. For example:

- 0T00: Present only at W1 in treated samples
- 0TTT: Present at W1, W2, and W3 in treated samples (persistent)
- 00TT: Appeared at W2 and persisted to W3
- 0C00: Present only at W1 in control samples

Structural variants were tracked across timepoints using coordinate-based matching with 1000bp tolerance for breakpoint positions, accommodating minor positional variability introduced by independent variant calling across samples. SVs on the same chromosome with the same SV type and breakpoints within tolerance were considered the same event. Point mutations were tracked using exact coordinate matching based on their unique identifiers (chromosome, position, reference allele, alternate allele). This enabled reconstruction of temporal trajectories for both structural variants and point mutations across the exposure period.

### 4.6 SV-Mutation Co-occurrence Analysis

To test whether mutations cluster near SV breakpoints, we performed systematic spatial enrichment analysis across multiple window sizes. Breakpoint coordinates were extracted from each SV: for inversions, deletions, duplications, and insertions, both 5’ and 3’ breakpoints were analyzed; for translocations, breakpoints on both source and partner chromosomes were included. Each unique breakpoint position was analyzed independently to avoid double-counting.

Mutations within specified distances from breakpoints (10bp, 25bp, 50bp, 100bp) were counted separately for radiation-induced (T-pattern) and control (C-pattern) mutations. For each chromosome, mutations were sorted by position and breakpoint-proximal mutations were identified using binary search for computational efficiency. A mutation was considered overlapping a window if its genomic coordinates intersected the breakpoint ± window region.

Enrichment was calculated as the ratio of observed to expected mutation counts, where expected counts were based on genome-wide mutation density:

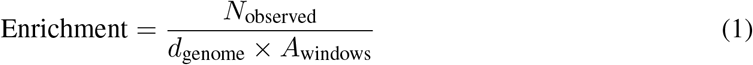

where *d*_genome_ is the genome-wide mutation density (mutations per bp) and *A*_windows_ is the total searchable area (number of unique breakpoints ×2 × window size).

The radiation-to-control enrichment ratio was calculated by dividing the enrichment for radiation-induced mutations by the enrichment for control mutations, providing a descriptive normalization to contextualize radiation-specific association against spontaneous mutation background. Statistical significance was assessed using Poisson tests comparing observed radiation mutation counts to expected counts based on genome-wide density. Enrichment analysis was performed in two stages: first across all SV breakpoints combined to assess general mutation clustering at structural variant junctions, and subsequently stratified by SV type to identify type-specific mutagenic coupling. SV type-specific enrichment was calculated separately for inversions, translocations, duplications, and deletions to identify type-specific mutagenic coupling. Distance-dependent decay was examined by comparing enrichment across increasing window sizes to determine the spatial extent of the mutagenic effect.

### 4.7 Repeat Element Analysis

Repeat element overlap was determined using AnnotSV annotations, which incorporate RepeatMasker classifications. SVs were categorized as having both breakpoints in repeats, one breakpoint in a repeat, or no repeat involvement. Interspersed repeat classes included LINE (Long Interspersed Nuclear Elements), Alu (a primate-specific SINE family derived from 7SL RNA), SINE (other Short Interspersed Nuclear Elements), LTR (Long Terminal Repeat retrotransposons), and DNA transposons; uncategorized interspersed elements were grouped as “Other”. Tandem and low-complexity repeats (e.g., `(CA)n`, alpha-satellite) were classified separately as “Simple” and excluded from interspersed-repeat involvement counts.

### 4.8 Temporal Concordance Analysis

To test whether spatially proximal INV-DBS pairs arise from shared mutagenic events, we performed temporal concordance analysis on co-occurring pairs identified within 10bp of inversion breakpoints. For each INV-DBS pair, we compared the inversion’s detection timepoint (W1, W2, or W3) against the DBS temporal pattern encoding (e.g., 0T00 for W1 only, 0TTT for W1+W2+W3). Pairs were matched on both radiation dose (A-E) and timepoint to ensure valid comparisons.

Temporal concordance was defined as the percentage of INV-DBS pairs where the inversion’s detection timepoint was present in the DBS pattern encoding. For example, an inversion detected at W2 (dose D) paired with a DBS showing pattern 00T0 (W2 only, dose D) would be concordant, while pairing with pattern 0T00 (W1 only, dose D) or 00TT (W2 and W3, dose D) would be discordant.

Expected concordance under the null hypothesis of independent events was calculated as the random probability of timepoint overlap given the empirical temporal distributions of inversions and DBS mutations. With three independent timepoints, random concordance was estimated at 33%. Statistical significance of observed concordance against this null expectation was assessed using a chi-square goodness-of-fit test at each timepoint and overall. Concordance was calculated overall and stratified by inversion size categories (small: *<*10kb, medium: 10kb–1Mb, large: 1–50Mb, mega: ≥50Mb).

### 4.9 Dose-Stratified Analysis and Functional Annotation

To assess dose-dependent effects on genomic instability, radiation exposures were stratified into two dose rate bins: Low (A-C: 0.20–0.40 mGy/hr) and High (D-E: 1.47–2.62 mGy/hr) [Roach et al., 2026]. INV-DBS co-occurrence analysis was performed separately for each dose bin to identify dose-specific genomic damage patterns.

Within each dose bin, INV-DBS pairs were identified with strict matching requirements: (1) spatial proximity (DBS within 10bp of inversion breakpoints), (2) dose concordance (same radiation dose level), and (3) temporal concordance (inversion detection timepoint present in DBS temporal pattern encoding). Only pairs satisfying all three criteria were retained for downstream analysis, ensuring identification of mutagenic events arising from shared repair processes rather than independent genomic alterations at the same locus. Genes overlapping inversions with temporally concordant DBS were extracted from AnnotSV annotations. Functional annotation was performed using the MyGene.info API (v3) [Xin et al., 2016] to retrieve gene names, functional summaries, Gene Ontology biological process terms, and pathway annotations (KEGG and Reactome). Constraint scores (pLI) were obtained from gnomAD v2.1.1 [Karczewski et al., 2020] to identify loss-of-function intolerant genes (pLI ≥0.9). Functional categorization was performed using keyword-based classification into six condensed categories: Signal Transduction, Gene Expression, Cell Structure & Adhesion, Cell Cycle & DNA Damage, Development & Differentiation, and Metabolism & Other. Category assignment was based on pattern matching across combined GO terms, pathway annotations, and gene function summaries. When genes matched multiple categories, assignment followed a priority hierarchy favoring Cell Cycle & DNA Damage, Signal Transduction, and Gene Expression over broader categories.

## 5 Workflow Infrastructure and Execution Substrate

We construct the spatio-temporal genomic analysis pipeline using a decoupled, asynchronous workflow orchestration stack to ensure reproducibility, scalability, and flexibility across heterogeneous computing systems. This infrastructure coordinates the execution of the multi-stage genomic variant annotation, mutational signature extraction, spatial correlation models, and statistical plotting modules and makes the pipeline portable across various execution platforms (Figure S5)

### 5.1 Workflow Orchestration

The complete analysis pipeline is decomposed into two coupled DAGs: the Mutation Analysis Pipeline (with 7 stages) and the Structural Variant (SV) Analysis Pipeline (with 14 stages), joined at the SV-mutation co-occurrence stage (4.6), where breakpoint tables and per-sample mutation tables are consumed jointly. RADICAL-AsyncFlow (RAF) [RADICAL-Cybertools Team, 2026a], an asynchronous workflow engine within the RADICAL Cybertools ecosystem [RADICAL-Cybertools Team, 2026b], orchestrates the execution and data dependencies of these workflows. In contrast to static workflow management systems that rely on compile-time dependency checking or file-pattern configurations, RADICAL-AsyncFlow utilizes Python’s native asynchronous event loop (`asyncio`) to dynamically resolve a Directed Acyclic Graph (DAG) at runtime. The engine dynamically schedules tasks as their direct dependencies (arguments and inputs) resolve.

### 5.2 Distributed Execution

The execution substrate integrated with the workflow management layer uses RHAPSODY, a high-performance multi-runtime middleware [Alsaadi et al., 2025] that decouples workflow composition from execution mechanics. Rather than replacing existing runtimes, RHAPSODY composes and coordinates them behind uniform abstractions for tasks, services, and resources, allowing heterogeneous workloads to share a single resource allocation. Each workflow selects an execution backend according to where it runs, ranging from local tests using the `Concurrent` backend to multi-node execution on leadership-class HPC platforms using a `Dask`-based (or `DragonHPC`-based) backend. This abstraction ensures that the execution of the same pipeline code is fully portable.

### 5.3 Enabled Scientific Capabilities

While workflow engines typically focus on engineering efficiencies, the integration of RADICAL-AsyncFlow and RHAPSODY introduces distinct scientific capabilities valuable for rigorous spatio-temporal genomic analysis.

1. *Dynamic Parameter-Space Sensitivity Analyses (Selection-Bias Mitigation)*. The asynchronous DAG runtime can schedule large numbers of parallel runs that vary analysis parameters, for example, the breakpoint-matching tolerance (`sv_tolerance`, 4.5) and the breakpoint-proximal window sizes (4.6), enabling systematic sweeps that test whether the reported couplings are threshold-independent rather than artifacts of specific cutoffs.
2. *Dynamic Runtime Adaptation and Conditional Analysis*. Biological data exploration is inherently iterative: the results of early stages frequently determine downstream analytical directions. RADICAL-AsyncFlow’s runtime event loop enables adaptive pipelines in which later stages are scheduled conditionally on upstream results, such as in our two-stage enrichment analysis (all breakpoints combined, then stratified by SV type) and the dose-stratified gene-impact analysis triggered by the INV-DBS signal. Such dynamic execution paths avoid manual, multi-step scripting interventions while maintaining strict data lineage.
3. *Co-scheduling of Heterogeneous Scientific Modalities*. Spatio-temporal genomics spans tasks with divergent computational profiles: (i) GPU-bound ML, (ii) memory-bound distributed analytics, and (iii) I/O bound metadata retrieval. By decoupling these tasks via RHAPSODY, the pipeline executes them within a unified resource allocation. A GPU worker can be provisioned exclusively for signature extraction when an optional GPU NMF mode is enabled (`SigProfilerExtractor` supports it), while high-throughput Dask workers handle distance decay calculations simultaneously. This prevents execution fragmentation, ensuring that cross-disciplinary analyses can be scaled in a unified, automated manner.
4. *Inter-lab Portability and Verification of Results*. Separating pipeline logic from hardware backends addresses the reproducibility challenge: the same execution code and parameters used for local developmental tests can be deployed unchanged on leadership-class HPC platforms. This ensures that findings are not tied to specific cluster schedulers, node configurations, or proprietary file systems, facilitating external verification and collaborative replication of the genomic instability models.

## Data Access

The whole-genome sequencing data generated for this study have been deposited in the NCBI Sequence Read Archive (SRA) under BioProject accession PRJNA1450698. All analysis code used to produce the results, figures, and tables presented in this manuscript is freely available at https://github.com/BNL-LUCID/rigi-analysis.

Previously published reference resources used in this study, including the GRCh38/hg38 reference genome and Mutect2 supporting files, were obtained from the Broad Institute public reference bundle (`gs://gcp-public-data-broad-references/hg38/v0/`).

## Acknowledgments

This research was funded by the Biological and Environmental Research program in the US DOE’s Office of Science under project B&R# KP1601017 and FWP#CC140. **K.C**.: Methodology, Software, Formal analysis, Investigation, Writing – original draft, Writing – review & editing. **C.C**.: Software (initial variant calling pipeline for point mutations), Data analysis, Writing – review & editing. **M.Ti**.: Software (workflow development), Methodology, Writing – original draft, Writing – review & editing. **R.W**.: Conceptualization, Investigation (experimental design and execution), Resources, Writing – review & editing. **S.F**.: Conceptualization, Investigation (experimental design and execution), Resources. **O.O.K**.: Software (workflow development), Methodology. **Y.Z**.: Conceptualization, Writing – review & editing. **M.Tu**.: Supervision (workflow development). **S.J**.: Supervision (workflow development). **D.S.S**.: Conceptualization, Investigation (experimental design), Resources, Supervision. **T.B**.: Conceptualization, Investigation (experimental design), Supervision. **B.J.Y**.: Supervision, Writing – review & editing. All authors reviewed and approved the final manuscript.

