## Supplementary material for "A Spatio-Temporal Analysis Framework for Characterizing Radiation-Induced Genomic Instability"

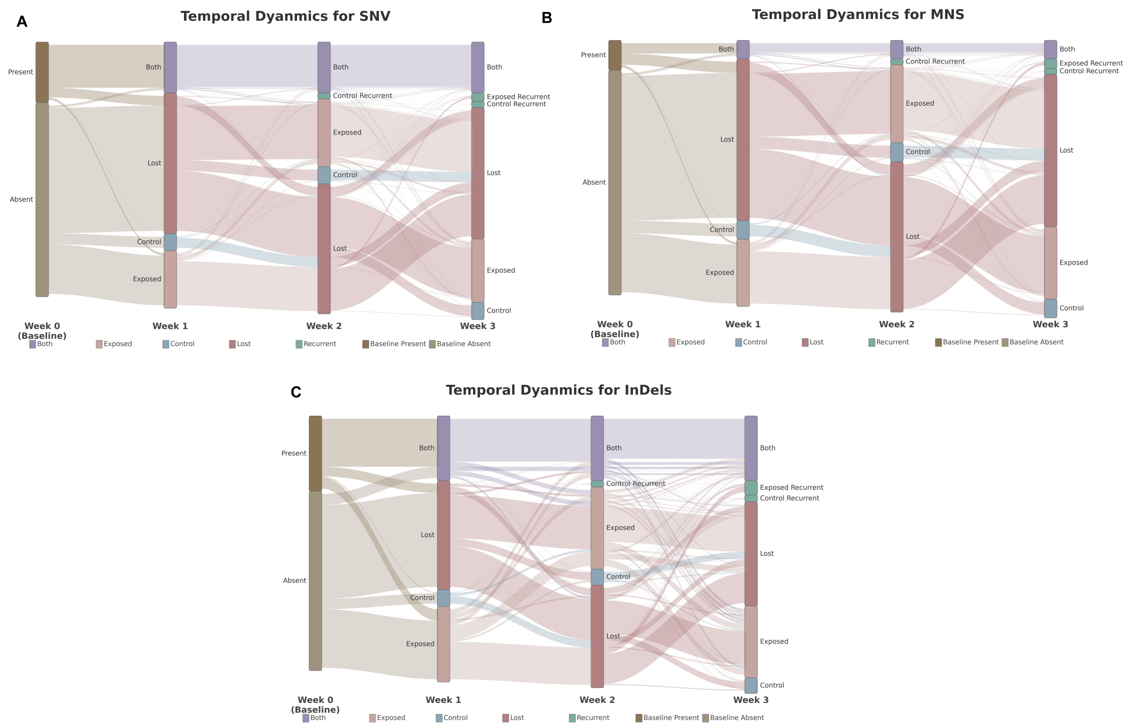


Figure S1: **Temporal dynamics of mutation accumulation for SNV, MNS, and InDel.** Sankey diagrams showing mutation flow across weekly timepoints (W0–W3), aggregated across all dose levels and chromosomes, for **(A)** single nucleotide variants (SNV), **(B)** multi-nucleotide substitutions (MNS), and **(C)** insertions/deletions (InDel). Flow width is proportional to mutation count; color coding follows Figure 2. SNV and InDel show a proportionally larger baseline-present pool with correspondingly stronger Present→Both flows persisting across all timepoints. MNS shows a smaller baseline-present contribution, with Exposed and Control flows representing a larger fraction of total mutation dynamics. Quantitative counts of radiation-specific temporal patterns across all mutation types are provided in Table S1.

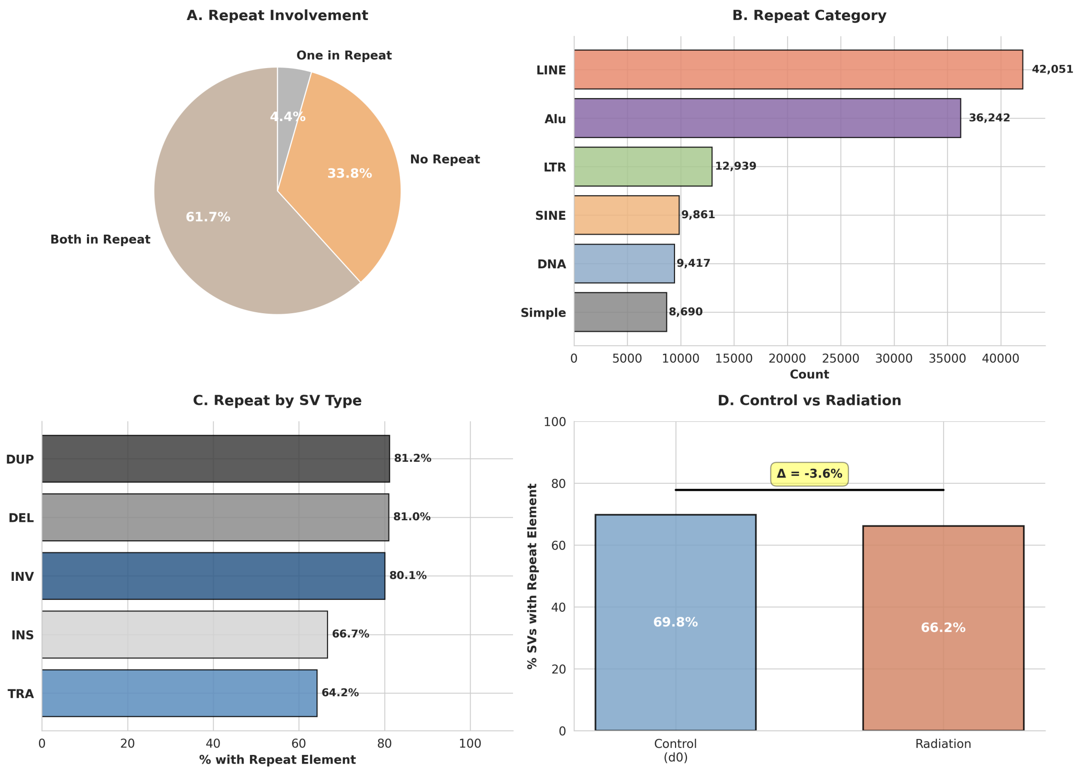


Figure S2: **Repetitive element involvement in structural variant breakpoints.** **(A)** Repeat element involvement at SV breakpoints. The majority of SVs have both breakpoints within interspersed repeat elements (61.7%), with 33.8% having no repeat involvement and 4.4% having one breakpoint in a repeat. **(B)** Distribution of repeat element categories at breakpoints. LINE elements predominate (42,051), followed by Alu (36,242), LTR (12,939), SINE (9,861), DNA transposons (9,417), and tandem/simple repeats (8,690). **(C)** Repeat involvement by SV type, defined as either breakpoint overlapping an interspersed repeat. Duplications (81.2%), deletions (81.0%), and inversions (80.1%) show high repeat association, while translocations (64.2%) and insertions (66.7%, n=3) show lower involvement. **(D)** Comparison of repeat involvement between control and radiation-induced SVs. Radiation SVs show slightly lower repeat involvement (66.2%) compared to control (69.8%, $\Delta = -3.6\%$), suggesting radiation may induce breakpoints in non-repetitive regions at modestly higher rates.

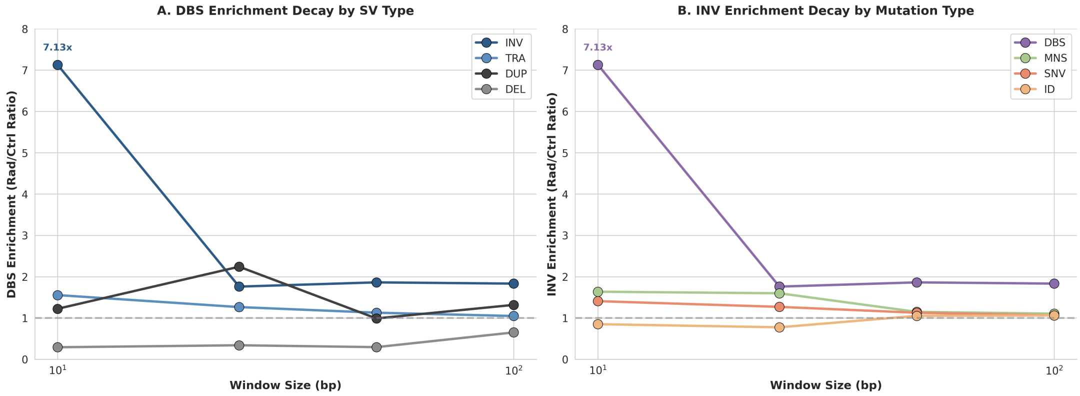


Figure S3: **Distance-dependent decay of mutation enrichment at structural variant breakpoints. (A)** DBS enrichment (radiation/control ratio) by SV type across increasing window sizes. INV-DBS coupling shows steep decay from 7.13× at 10bp to ∼1.9× at 100bp, indicating tight spatial localization of mutagenesis at inversion breakpoints. TRA, DUP, and DEL show minimal or no enrichment across all distances. **(B)** Mutation type enrichment at INV breakpoints across window sizes. DBS shows strong distance-dependent decay (7.13× to ∼1.8×), while MNS, SNV, and InDel remain near baseline (0.8–1.6×) regardless of distance. Dashed lines indicate expected baseline (1.0×). The rapid decay pattern suggests DBS arise from error-prone repair directly at INV breakpoint junctions rather than from regional mutagenic effects.

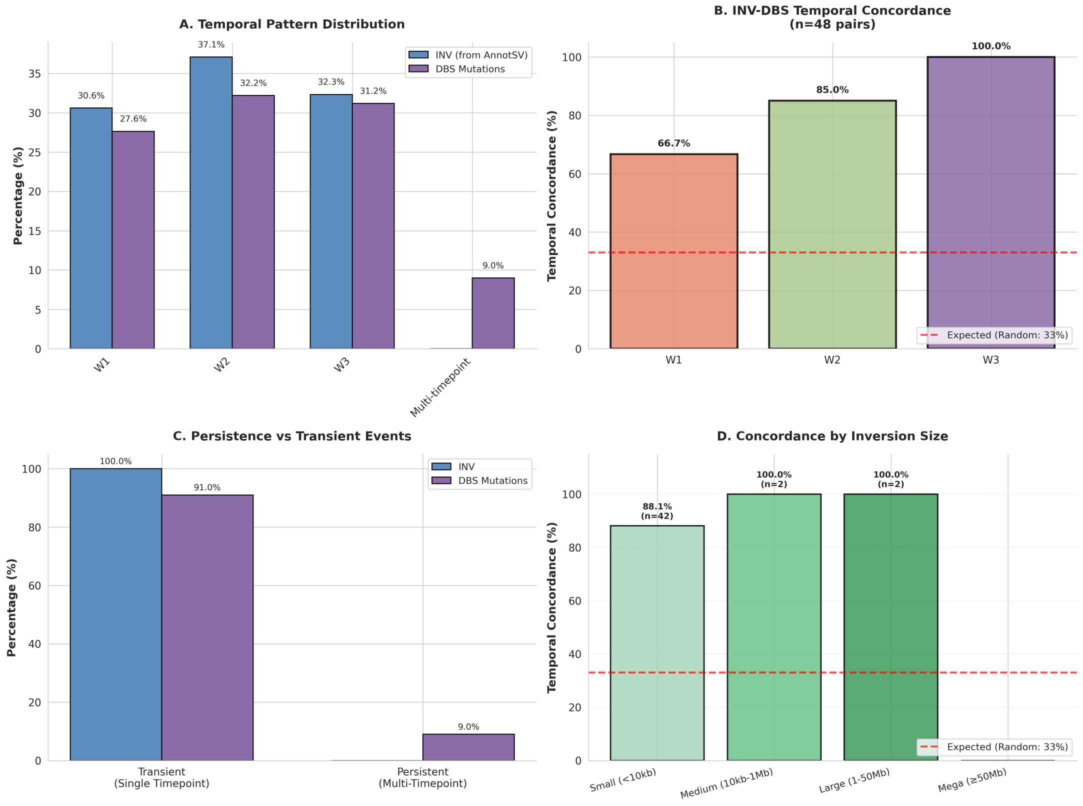


Figure S4: **Temporal dynamics of inversion-DBS co-occurrence.** **(A)** Temporal distribution of inversions (blue) and DBS mutations (purple). Both show similar accumulation patterns across timepoints (W1: 30.6% vs 27.6%, W2: 37.1% vs 32.2%, W3: 32.3% vs 31.2%). DBS show 9.0% multi-timepoint persistence while inversions are exclusively single-timepoint. **(B)** Temporal concordance between spatially co-occurring INV-DBS pairs (n=48 pairs within 10bp, matched by dose). Concordance increases over time (W1: 66.7%, W2: 85.0%, W3: 100%), all exceeding the 33% random expectation, indicating INV and nearby DBS likely arise from shared mutagenic processes rather than independent events at the same locus. **(C)** Transient versus persistent events. Inversions are exclusively transient (100% single-timepoint), while DBS show modest persistence (9.0% multi-timepoint), reflecting differential cellular quality control. **(D)** Concordance by inversion size. Concordance was high for small (<10kb, n=42; 88.1%), medium (10kb–1Mb, n=2; 100%), and large (1–50Mb, n=2; 100%) inversions, while mega-inversions (≥50Mb, n=2) showed no temporally concordant DBS pairs (0%), suggesting different mutagenic dynamics for the largest rearrangements.

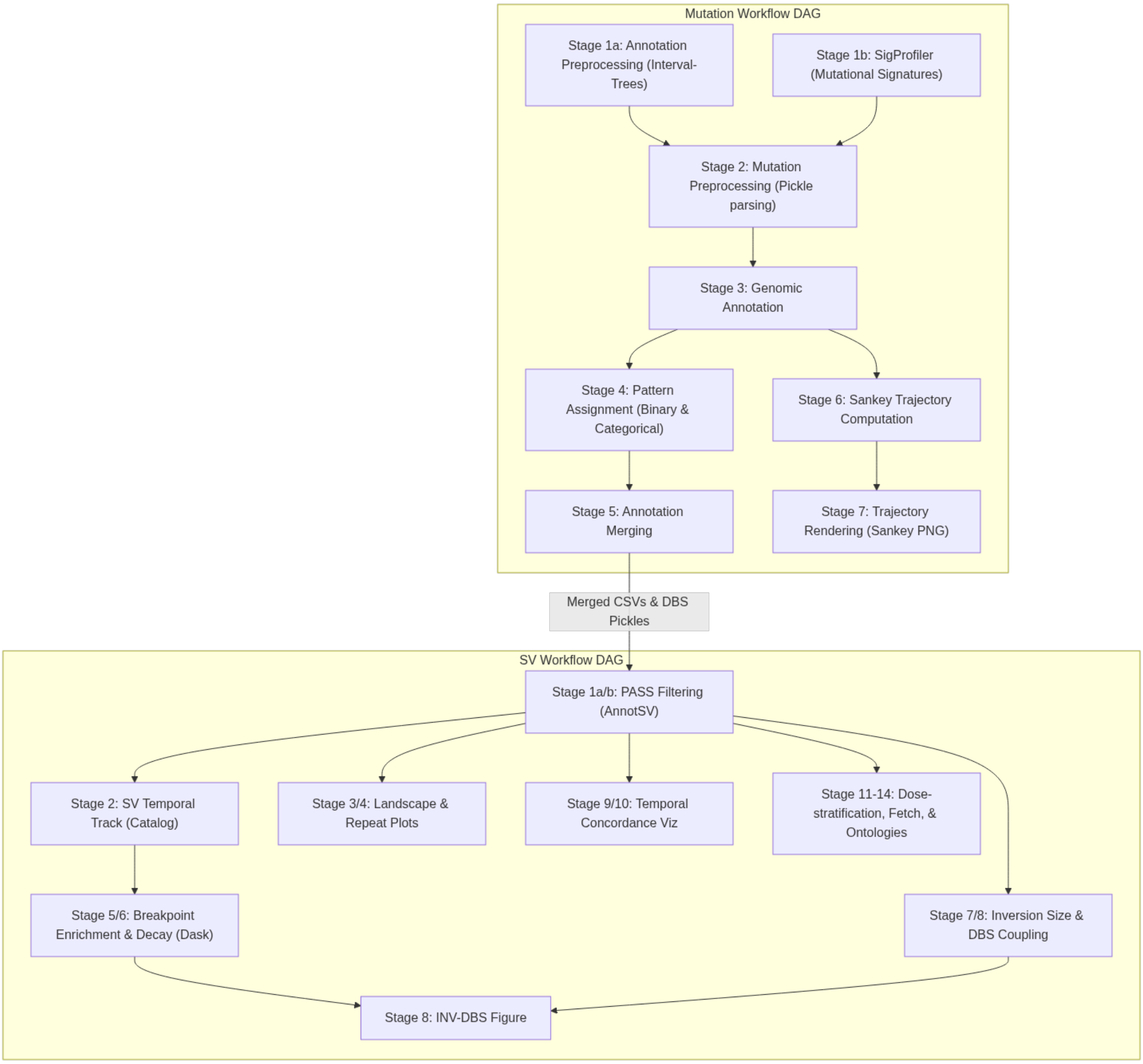


Figure S5: **Workflow orchestration of the spatio-temporal analysis pipeline.** Directed acyclic graph (DAG) representation of the two coupled workflows that implement the analysis, orchestrated by RADICAL-AsyncFlow within the RADICAL-Cybertools ecosystem. The Mutation Workflow (top) processes point mutations through annotation preprocessing, SigProfiler signature profiling, genomic annotation, temporal pattern assignment, and annotation merging, with a parallel branch computing and rendering the Sankey trajectories. The Structural Variant Workflow (bottom) begins with PASS-filtered, AnnotSV-annotated SVs and branches into temporal tracking, landscape and repeat characterization, breakpoint-proximal enrichment and distance decay, temporal concordance, inversion-size and DBS coupling, and dose-stratified functional annotation. The two workflows are joined at the SV-mutation co-occurrence stage, where merged mutation and breakpoint tables are consumed together to generate the INV-DBS results. Boxes denote analysis stages and arrows denote data dependencies resolved at runtime; the modular structure allows the same pipeline to execute unchanged across local and HPC backends via RHAPSODY.

### Supplementary Tables

Table S1: **Radiation-induced mutation counts by temporal pattern and mutation type.** Counts represent unique mutations detected exclusively in radiation-exposed samples (T-pattern), summed across all dose levels (A–E) and all chromosomes. Pattern notation follows the four-position encoding described in Methods (W0, W1, W2, W3), where `0` = absent and `T` = exposed only. Single-timepoint patterns reflect transient mutagenic events; multi-timepoint patterns reflect clonal persistence; the recurrent pattern (`0T0T`) reflects re-emergence after clearance.

| Pattern | Timepoints detected | SNV | DBS | MNS | InDel |
|---|---|---|---|---|---|
| *Single-timepoint* | | | | | |
| `0T00` | W1 only | 5,086,175 | 79,285 | 448,130 | 716,513 |
| `00T0` | W2 only | 6,176,589 | 94,527 | 533,752 | 808,200 |
| `000T` | W3 only | 5,975,176 | 90,835 | 518,469 | 785,579 |
| *Multi-timepoint* | | | | | |
| `0TT0` | W1 + W2 | 113,684 | 2,444 | 9,999 | 59,901 |
| `00TT` | W2 + W3 | 127,584 | 2,658 | 10,946 | 64,235 |
| `0T0T` | W1 + W3 (recurrent) | 113,375 | 2,505 | 10,269 | 60,562 |
| *Persistent* | | | | | |
| `0TTT` | W1 + W2 + W3 (persistent) | 29,384 | 595 | 1,918 | 19,746 |
| | **Total** | **17,621,967** | **272,849** | **1,533,483** | **2,514,736** |

Table S2: Family-level repeat distribution at SV breakpoints in control versus radiation-induced events. Counts represent breakpoint occurrences (LEFT + RIGHT breakpoints combined) classified into interspersed repeat families and tandem/simple repeats. Percentages are calculated as a fraction of total breakpoints within each group. $\Delta$ denotes the difference in percentage points (Radiation − Control).

| Family | Control | Control % | Radiation | Radiation % | $\Delta$ (pp) |
|---|---|---|---|---|---|
| LINE | 4,994 | 23.0% | 42,051 | 23.5% | +0.6 |
| Alu | 5,578 | 25.7% | 36,242 | 20.3% | −5.4 |
| SINE | 1,094 | 5.0% | 9,861 | 5.5% | +0.5 |
| LTR | 1,582 | 7.3% | 12,939 | 7.2% | $\sim 0$ |
| DNA | 1,037 | 4.8% | 9,417 | 5.3% | +0.5 |
| Simple | 1,141 | 5.2% | 8,690 | 4.9% | −0.4 |
| Other | 431 | 2.0% | 3,725 | 2.1% | +0.1 |
| None | 5,877 | 27.0% | 55,673 | 31.2% | +4.1 |
| **Total** | **21,734** | | **178,598** | | |

Table S3: **Mutation enrichment at structural variant breakpoints across window sizes and mutation types.** Enrichment calculated as the ratio of observed to expected mutation counts, where expected counts are based on genome-wide mutation density (mutations per bp $\times$ total searchable window area). Window area $= 2 \times$ window size $\times$ 77,951 unique breakpoints. $P$-values from one-sided Poisson test. $^{***}P < 0.001$; $^{**}P < 0.01$; $^{*}P < 0.05$; n.s., not significant.

| Mutation Type | Window (bp) | Class | Observed | Expected | Enrichment | $P$-value | |
|---|---|---|---|---|---|---|---|
| DBS | 10 | Radiation | 293 | 189 | 1.55 | $1.65 \times 10^{-12}$ | *** |
| | | Control | 184 | 193 | 0.96 | $7.41 \times 10^{-1}$ | n.s. |
| | 25 | Radiation | 547 | 473 | 1.16 | $4.59 \times 10^{-4}$ | *** |
| | | Control | 441 | 481 | 0.92 | $9.70 \times 10^{-1}$ | n.s. |
| | 50 | Radiation | 908 | 946 | 0.96 | $8.93 \times 10^{-1}$ | n.s. |
| | | Control | 826 | 963 | 0.86 | 1.00 | n.s. |
| | 100 | Radiation | 1,706 | 1,891 | 0.90 | 1.00 | n.s. |
| | | Control | 1,569 | 1,926 | 0.81 | 1.00 | n.s. |
| SNV | 10 | Radiation | 13,895 | 11,175 | 1.24 | $< 1.0 \times 10^{-16}$ | *** |
| | | Control | 13,695 | 11,998 | 1.14 | $< 1.0 \times 10^{-16}$ | *** |
| | 25 | Radiation | 31,124 | 27,937 | 1.11 | $< 1.0 \times 10^{-16}$ | *** |
| | | Control | 30,842 | 29,996 | 1.03 | $5.89 \times 10^{-7}$ | *** |
| | 50 | Radiation | 59,020 | 55,875 | 1.06 | $< 1.0 \times 10^{-16}$ | *** |
| | | Control | 59,666 | 59,992 | 0.99 | $9.09 \times 10^{-1}$ | n.s. |
| | 100 | Radiation | 115,983 | 111,749 | 1.04 | $< 1.0 \times 10^{-16}$ | *** |
| | | Control | 120,314 | 119,984 | 1.00 | $1.71 \times 10^{-1}$ | n.s. |
| MNS | 10 | Radiation | 1,284 | 984 | 1.30 | $< 1.0 \times 10^{-16}$ | *** |
| | | Control | 1,197 | 1,032 | 1.16 | $3.12 \times 10^{-7}$ | *** |
| | 25 | Radiation | 2,562 | 2,461 | 1.04 | $2.18 \times 10^{-2}$ | * |
| | | Control | 2,341 | 2,581 | 0.91 | 1.00 | n.s. |
| | 50 | Radiation | 4,518 | 4,922 | 0.92 | 1.00 | n.s. |
| | | Control | 4,647 | 5,162 | 0.90 | 1.00 | n.s. |
| | 100 | Radiation | 9,164 | 9,844 | 0.93 | 1.00 | n.s. |
| | | Control | 9,189 | 10,325 | 0.89 | 1.00 | n.s. |
| InDel | 10 | Radiation | 2,970 | 2,547 | 1.17 | $2.22 \times 10^{-16}$ | *** |
| | | Control | 3,022 | 2,601 | 1.16 | $4.44 \times 10^{-16}$ | *** |
| | 25 | Radiation | 6,793 | 6,368 | 1.07 | $7.20 \times 10^{-8}$ | *** |
| | | Control | 6,953 | 6,503 | 1.07 | $1.77 \times 10^{-8}$ | *** |
| | 50 | Radiation | 15,112 | 12,737 | 1.19 | $< 1.0 \times 10^{-16}$ | *** |
| | | Control | 15,218 | 13,006 | 1.17 | $< 1.0 \times 10^{-16}$ | *** |
| | 100 | Radiation | 28,100 | 25,473 | 1.10 | $< 1.0 \times 10^{-16}$ | *** |
| | | Control | 29,324 | 26,012 | 1.13 | $< 1.0 \times 10^{-16}$ | *** |

Table S4: **High-constraint genes (pLI $\geq$ 0.9) affected by temporally concordant INV-DBS pairs.**

| Gene | Chr | pLI | Category | Function |
|---|---|---|---|---|
| **HIGH DOSE (15 genes)** | | | | |
| WAC | 10 | 1.00 | Cell Cycle & DNA Damage | Regulates cell-cycle checkpoint activation in response to DNA damage |
| CUL2 | 10 | 1.00 | Cell Cycle & DNA Damage | E3 ubiquitin-protein ligase complex component |
| RET | 10 | 1.00 | Signal Transduction | Proto-oncogene tyrosine-protein kinase receptor |
| RASGEF1A | 10 | 0.97 | Signal Transduction | Guanyl-nucleotide exchange factor; involved in cell migration |
| MAP3K8 | 10 | 0.97 | Signal Transduction | MAPK/ERK pathway activation; pro-inflammatory cytokine production |
| EPC1 | 10 | 1.00 | Gene Expression | NuA4 HAT complex; DNA double-strand break repair via HR |
| ZEB1 | 10 | 0.97 | Gene Expression | Zinc finger transcription factor; transcriptional repressor |
| PATL1 | 11 | 0.96 | Gene Expression | mRNA deadenylation-dependent decapping and degradation |
| KIF5B | 10 | 0.99 | Cell Structure & Adhesion | Centrosome and nuclear positioning during mitosis |
| ITGB1 | 10 | 0.98 | Cell Structure & Adhesion | Cell adhesion during telophase; required for cytokinesis |
| PARD3 | 10 | 0.96 | Cell Structure & Adhesion | Asymmetrical cell division and cell polarization |
| RAB18 | 10 | 0.91 | Cell Structure & Adhesion | Intracellular membrane trafficking regulation |
| NRP1 | 10 | 0.99 | Development & Differentiation | Cardiovascular development, angiogenesis, neuronal circuits |
| SYT2 | 1 | 0.98 | Development & Differentiation | Dendrite formation by melanocytes |
| MTPAP | 10 | 1.00 | Metabolism & Other | Mitochondrial RNA poly(A) tail polymerase |
| **LOW DOSE (1 gene)** | | | | |
| MKLN1 | 7 | 1.00 | Cell Structure & Adhesion | Cell spreading and cytoskeletal responses to ECM |
